# CONE: COntext-specific Network Embedding via Contextualized Graph Attention

**DOI:** 10.1101/2023.10.21.563390

**Authors:** Renming Liu, Hao Yuan, Kayla A Johnson, Arjun Krishnan

## Abstract

Human gene interaction networks, commonly known as interactomes, encode genes’ functional relationships, which are invaluable knowledge for translational medical research and the mechanistic understanding of complex human diseases. Meanwhile, the advancement of network embedding techniques has inspired recent efforts to identify novel human disease-associated genes using canonical interactome embeddings. However, one pivotal challenge that persists stems from the fact that many complex diseases manifest in specific biological contexts, such as tissues or cell types, and many existing interactomes do not encapsulate such information. Here, we propose CONE^3^, a versatile approach to generate context-specific embeddings from a context-free interactome. The core component of CONE consists of a graph attention network with contextual conditioning, and it is trained in a noise contrastive fashion using contextualized interactome random walks localized around contextual genes. We demonstrate the strong performance of CONE embeddings in identifying disease-associated genes when using known associated biological contexts to the diseases. Furthermore, our approach offers insights into understanding the biological contexts associated with human diseases.

## 1 Introduction

The proper operation of cells depends on the precise coordination and interaction of biological entities, such as genes, RNA, and proteins. As a result, complex human diseases are the ramifications of perturbations to groups of genes that give rise to pathological states [1, 2]. Leveraging this interdependence among biological entities, network-based methods have shown great promises in unveiling human genes’ function [3] and their associated diseases [4, 5]. Recent approaches achieved this by training machine learning models using network embeddings extracted from the input gene network [3, 6]. However, a crucial limitation remains as many biological network embedding methods do not consider the differences induced by various biological contexts.

The interacting relationships among genes vary across biological contexts, such as tissues, cell types, or disease states. Many human genes operate in a tissue-dependent manner. For example, *DMD* is preferably expressed in muscle [7]. The heterogeneity of the specific set of genes expressed by a particular biological context contributes significantly to the phenotypic diversity within an organism, aiding the complex, specialized functions required for its survival [8]. Consequently, the dysfunction of genes ultimately leads to diseases manifesting only in specific tissues. For example, Mendelian disorders show clear tissue-specific manifestation and complex diseases have a strong tendency of tissue selectivity, such as neurological disorders and cardiovascular diseases [9]. However, tissue-disease associations are not always straightforward. Apart from the primary affected tissues, diseases may affect seemingly unrelated tissues. One prime example is the high risk of gastrointestinal tract dysfunction observed in patients of Parkinson’s disease, a neurological disorder primarily centered in the brain [10]. The cryptic connections between diseases may be partially explained by shared underlying mechanisms among them, which could be well characterized by networks [11].

Numerous functional genomics projects generate data comprising diverse types, qualities, and scopes of genes or molecules [12–14]. To obtain a high-quality and comprehensive network embedding, several methods are developed to infer a joint network representation by integrating multiple networks [15–19]. However, a drawback of network integration methods is that the integration process can eliminate context-specific information in each input network, resulting in a context-naive network. In other words, it may assume the same molecular interactions in the kidney and brain, whereas in reality, interactions are tissue-specific. To predict a range of tissue-specific gene functions or gene-disease relationships, we must integrate context-specific information into the network. Furthermore, state-of-the-art data integration approaches using graph neural networks [16] may not scale well to the number or size of the networks. Therefore, we require a scalable method that can handle networks of varying sizes and numbers.

### Contributions

Here, we address the critical need for a versatile and scalable method for generating biological context-specific network embeddings. We summarize our main contributions as follows.

1. We propose CONE, a versatile contextual network embedding method that takes context definitions in the form of node sets.
2. The proposed method operates on a shared graph attention network across all contexts, which is contextualized by conditioning on the raw embeddings. This results in a model that scales practically independent to the number of contexts.
3. Through a series of experiments, we demonstrate the value of injecting various biological contexts to improve disease gene prioritization.

## 2 Related work

A few studies have explored the idea of contextualizing biological network embeddings using contexts such as tissue or cell type specificity. Notably, OhmNet [20] pioneered the tissue-specific gene interaction network embedding by leveraging the hierarchical relationships between different tissue levels and genes. OhmNet learns a multi-layer embedding and operates on the idea that closely related tissues, or layers, should have similar embeddings. However, the original OhmNet method requires a highly specific construction of the hierarchical multi-layer tissue-specific networks, making it hard to extend to broader biological contexts readily. More recently, PINNACLE [21] further expanded the biological contexts into finer-grain definitions based on cell types using cell-type-expressed genes constructed from the Tabula Sapiens single cell atlas [22]. Furthermore, PINNACLE learns context-specific graph attention modules with independent parameters per context.

## 3 Preliminaries

### 3.1 Network Embedding via Sampling

Let *G* = (*V, E, w*) be a weighted undirected graph, where the edge weight function *w* : *V × V →* ℝ, and denote its corresponding adjacency matrix by **A** *∈* ℝ^|*V* |*×*|*V* |^. The goal of a graph embedding method aims to find the mapping *f* : *V→* ℝ^*d*^ that maps each node *v ∈ V* to a *d*-dimensional embedding space by minimizing the following objective function.

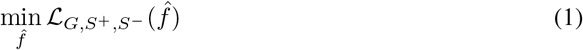

where *S*^+^ and *S*^*−*^ are positive and negative edge sampling functions. Particularly, the Singular Value Decomposition (SVD) can be viewed as the optimization of equation 1, where *S*^+,*−*^ are both uniform sampler on all pairwise entries in the adjacency matrix, *f* maps to the left (*f*_*L*_) and right (*f*_*R*_) embedding representations [23]. The loss function is the squared error between the inner product of the left and right embeddings of the two nodes and the edge weight between them in the graph:

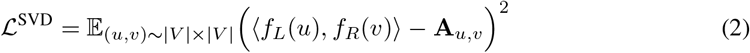

### Random Walk Sampling

Random walk on graphs have been studied extensively, with many applications spanning social network analysis, information retrieval, and so on. In our framework, the random walk procedure can be seen as the node-pair sampling function. For instance, *node2vec* [24] with negative sampling can be reformulated in a noise contrastive fashion [25] as:

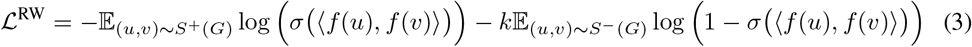

where *σ*is the sigmoid function, *k* is the number of negative samples, and the positive sampling is achieved by a sliding window over a second-order biased random walk [24]. We refer to the above as the *random walk loss*.

### 3.2 Graph Attention Neural Network (GAT)

Graph neural network (GNN) is a special type of neural network architecture that operates on the underlying graph structure. It does so by iteratively aggregating information from each node’s neighborhood and transforming the aggregated representations [26, 27]. Particularly, GAT [28, 29] uses an attention mechanism to weight each node’s neighborhood for aggregation, and the (preactivation and pre-normalization) layer updating rule is written as follows.

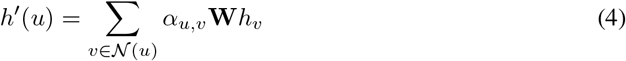

where *α*_*u,v*_ is the attention score, **W** *∈* ℝ^*d×d*^ is a learnable linear transformation. In practice, we use the v2 corrected attention proposed in [29]:

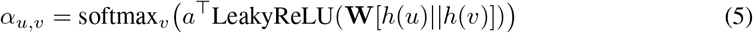

## 4 Our Method

We are interested in learning a *collection* of network embeddings, each specific to a biological context. For example, we can use heart-specific gene embeddings to unravel more tissue-specific genes related to cardiovascular diseases. Contextualizing gene embeddings to biological contexts this way allows us to unveil nuanced relationships between diseases and biological contexts, such as tissues, cell types, and other diseases or traits. The full pipeline of our approach is depicted in Figure 1.

**Figure 1:**
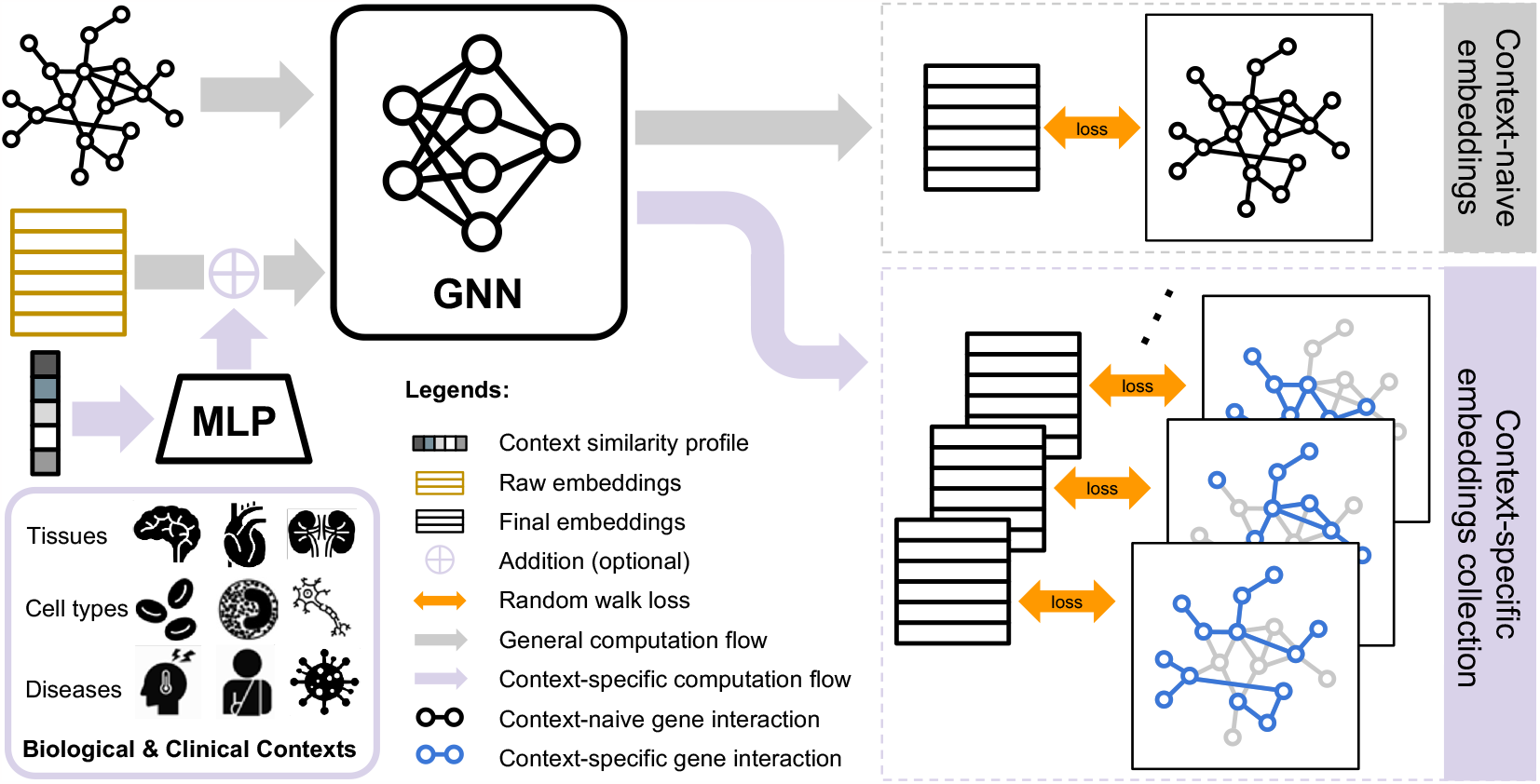
Overview of CONE embedding collection training and inference.

From a high level, CONE contains two main components, including (1) a GNN decoder and (2) an MLP context encoder. The GNN decoder converts the raw, learnable, node embeddings into the final embeddings. On the other hand, the MLP context encoder projects the context-specific similarity profile that describes the relationships among different contexts (Section 4.2) into a condition embedding. When added with the raw embeddings, the condition embedding serves as a high-level contextual semantics, similar to the widely-used positional encodings in Transformer models [30]. The embeddings are trained using the losses based on the random walk on the context-specific subgraphs. We employ a straightforward approach to define a context-specific subgraph as the subgraph induced by the genes relevant to that context. Next, we formally describe our approach.

### 4.1 Contextualized network embeddings

Let *C* = *{C*_*i*_*}*_*i∈*1,…,*n*_*c* be a collection of *n*_*c*_ contexts, where each context *C*_*i*_ *⊂ V* is a subset of nodes that defines the local context. We aim to learn a collection of embedding functions ℱ = *{f*_*C*_*}* by

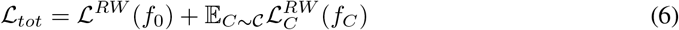

where *f*_0_ is the context-naive embedding that is optimized against the whole network *G*, and *f*_*C*_ is the context-specific embedding that is optimized against the contextual random walk loss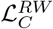 that samples random walks on the subgraph induced on the context set *C*, that is, *G*(*C*) = (*C, {* (*u, v*) *∈ E* : *u, v ∈ C}, w*). Equation 6 aims to simultaneously optimize for the global and local contextualized representation of all the nodes *v* in the network.

A naive attempt for obtaining the contextualized embeddings in equation 6 would be to learn independent *f*_*C*_ on the corresponding contextual graph *G*(*C*). However, the resulting context-specific embeddings may completely lose the global information of the graph, since each *f*_*C*_ operates independently. We provide empirical evidence for this in Section B.2.

### 4.2 Contextualized GAT

To address the above-mentioned problem, we propose to learn a *shared embedding encoding model* using GAT, and contextualize different embeddings by conditioning on the raw embedding matrix.

Let *g*_*θ*_ :ℝ^|*V*|*×d*^*→*ℝ^|*V* |*×d*^ be a GAT network parameterized by *θ*, and **Z** *∈* ℝ^|*V* |*×d*^ the raw embedding matrix that is randomly initialized. Drawing parallels from recent work on conditional generation [31], we view context-specific embeddings as generation conditioned on a specific context, and propose to compute the contextualized embedding *f*_*C*_ as

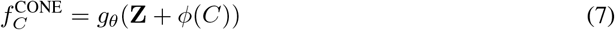

where *ϕ*(*C*) *∈* ℝ^1*×d*^ is the context condition embedding that defines the context *C*. The context-naive embeddings are computed by passing the raw embedding alone through the GAT encoder: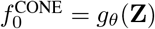. Finally, to form the full context-specific embedding for downstream evaluation, we concatenate it with the context-naive embedding and then project it down to *d*-dimension via PCA.

### Context condition embedding

The context condition embedding serves to provide low level semantics about each context, and two contexts with highly overlapping sets of nodes should have similar condition embeddings. To that end, we design an approach to encode condition embedding using the context similarity matrix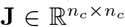 constructed by taking the Jaccard index between all pairwise contexts,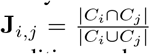. Finally, we use a two-layer Multi-Layer Perceptron (MLP) to project **J** into the condition embeddings, thus for each context *C*_*i*_, its corresponding condition embedding is computed as *ϕ*(*C*_*i*_) = MLP(**J**)_[*i*,:]_.

### 4.3 Training CONE

The loss function defined in equation 6 is implemented in practice by alternating between the context-naive random walk loss ℒ ^*RW*^ (*f*_0_), and the context-specific random walk loss ℒ ^*RW*^ (*f*_*C*_) for a randomly drawn context. We train the model for 120 epochs using the AdamW [32] optimizer with a constant learning rate of 0.001 and a weight decay of 0.01. Hyperparameter selection details can be found in Appendix C.2.

### 4.4 Complexity analysis

As the main module of CONE, GAT has the computational complexity of *O* (|*V*| *d*^2^ + |*E*| *d*) [28, 29]. In addition, the context condition embedding encoder *ϕ* scales linearly with respect to the number of conditions as *O* (*n*_*c*_*d*). However, in practice, since *n*_*c*_ *≪*|*E*|, the computational complexity of CONE should be equivalent to that of a single GAT network. This effective constant scaling with respect to the number of contexts is in stark contrast with the recently proposed method PINNACLE, which scales linearly with respect to the number of contexts due to the implementation of independent GAT module for each context. We provide empirical evidence for the scalability of CONE in Appendix A.

## 5 Experiments

### 5.1 Setup

We devise diverse biomedical tasks to evaluate the capability of CONE against baseline methods to prioritize genes in the gene interaction network. These tasks are binary classification tasks, where the goal is to identify human genes that are related to certain diseases using the gene network embeddings generated by the models. We conduct our main analysis using the PINPPI network, which is a combined network using BioGRID [33], Menche [34], and HuRI [35], provided by the PINNACLE [21] paper. For gene label information, we collect the two therapeutic target tasks (RA and IBD) from PINNACLE. Furthemore, we compile a comprehensive collection of disease-gene annotations from DisGeNET [36], following the processing steps detailed in [3]. After filtering out diseases with less than ten positive genes intersecting with the PINPPI network, the final DisGeNET benchmark contains 167 diverse human diseases.

For each DisGeNET disease gene prioritization tasks, we randomly split the positive and negative genes into 6/2/2 train validation test sets. The final prediction performance are reported as the average test scores across five different random splits. For RA and IBD, we use the pre-defined train test split given by PINNACLE. Detailed dataset statistics and processing notes can be found in Appenxi C.1

### Baselines

*node2vec* is a strong baseline method for network embedding-based gene prioritization method with superior performance on various benchmarks [37]. We also include embeddings generated by a two-layer GAT (v2) network [28, 29] trained in a standard graph autoencoder style [38] as a more direct baseline against CONE. Moreover, BIONIC [16] and Gemini [19] are two recent approaches that learn an integrated embedding across a collection of networks. We use them to test if embedding multiple context-specific subgraphs together gives an advantage over embedding a single context-naive network. All baselines and the CONE embeddings are evaluated in an *unsupervised* setting, where an *ℓ*_2_ regularized logistic regression model is trained for each task using the embeddings that are learned without accessing any label information.

For context-specific network embeddings, we consider a recently proposed method, PINNACLE [21], which learns separate GAT modules for each context. We directly use the context-specific embeddings provided by the paper ^4^ to reanalyze the performance under our fair setting. We point out that PINNACLE context-specific embeddings differ slightly from CONE in that PINNACLE only generates embedding for the context-specific nodes. Conversely, CONE generates embeddings for all nodes regardless of if they are specific to the context. This enables us to evaluate all context-specific embeddings fairly across diverse disease gene prioritization tasks. Due to this limitation of PINNACLE, we exclude it from the main disease gene prioritization benchmark (***RQ1***). We set the embedding dimensions to 128 for all models.

### Context-specific gene sets

We primarily consider tissue-specificity for contextualizing the network embeddings. This leads to a natural understanding of the downstream disease gene prediction performance based on the association between tissues (contexts) and diseases (tasks). One widely adopted way of defining tissue- or cell type-specific genes is by *differential gene expression* [39], where genes that express significantly more in one context than others are considered relevant to the given context. Following this, we first obtain tissue-specific gene expression from the GTEx project [40], and then extract tissue-specific genes by taking genes with z-scores greater than one across tissues. In addition to tissue-specific genes, we also showcase CONE using other biological contexts, including cell types and diseases. The specific constructions for those context-specific gene sets and their statistics can be found in Appendix C.1.

### Metrics

We use the log_2_ fold-change of the average-precision over the prior (APOP) as the main metric for evaluating disease gene prioritization performance [3]. The area under the precision recall curve, which is closely related to the average precision, is a better choice for evaluating tasks with severe class imbalance [41]. The prior division, on the other hand, corrects for the expectation of the random guesser performance on different tasks with different number of positive examples. For the RA and IBD therapeutic target prediction tasks, we follow PINNACLE and report the average precision and recall at rank five (APR@5) in addition to APOP.

### 5.2 Results and Discussions

#### RQ1. Can context-specific embedding improve disease gene prediction performance?

We first observe that, overall, picking the best context for each disease achieves noticeable performance improvement over the context-naive CONE embeddings, as indicated by the good performance of CONE (best) in Figure 2. Moreover, the advantage of using context-specific embedding is most apparent when the number of positive genes available for the disease is lacking, which might be attributable to the fact that diseases with more associated genes are more likely to contain more ubiquitous and less context-specific genes, and consequently reducing the effectiveness of using context-specific embeddings. In all cases, we note that either the context-naive or the context-specific CONE embedding consistently match the performance achieved by the *node2vec* baseline.

**Figure 2:**
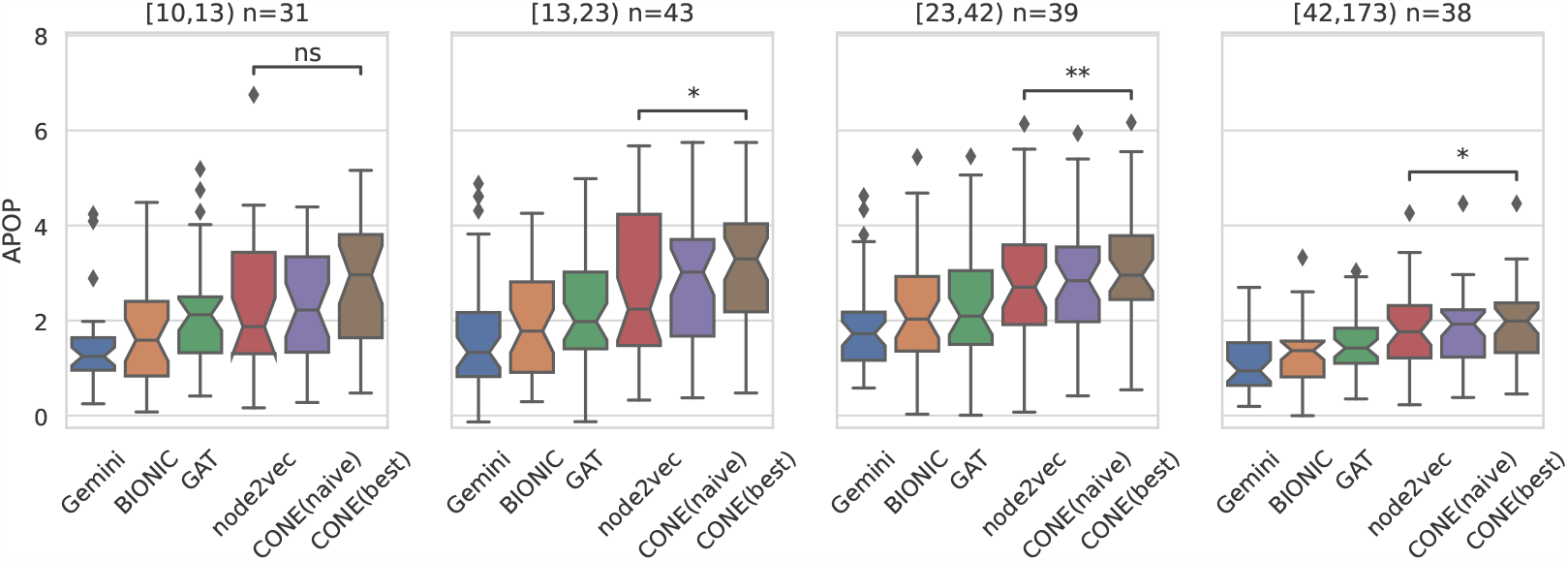
DisGeNET disease gene predictions performance comparison between node2vec and CONE embeddings. Each point in the box plot corresponds to the prediction test performance of a disease averaged across five random splits. Different panels show groups of diseases with different number of positive genes. For example, the left-most panel contains 31 diseases with at least 10 but less than 13 positive genes. ns, *, and ** indicate the significance level of the paired Wilcoxon test between the baseline *node2vec* and CONE (ns: not significant, *: 0.01*<* p-value*≤* 0.05, **: p-value < 0.01).

#### RQ2. Are the top performing context biologically relevant?

Despite the performance improvement due to the CONE contextualized embeddings, it is still unclear whether the biological contexts that led to good performance on a particular disease are associated with that disease. To address this question, we manually inspect six diseases where the connection between tissue and disease manifestation appears readily evident, as shown in Table 1. We hypothesize that the CONE embeddings derived from the disease-related tissue should have a higher APOP compared to the context-naive CONE or node2vec embeddings. Indeed, many of the top contexts found by CONE are biologically meaningful, whether as one of the main affected tissues or a related tissue. For example, the top-performing contexts for subvalvular aortic stenosis and familial bicuspid aortic valve, both diseases in which there is a problem with the aortic valve, which is the valve of an artery in the heart, included artery for subvalvular aortic stenosis and heart for familial bicuspid aortic valve. More subtle top-performing but biologically-relevant contexts are small intestine and adipose for hypochromic microcytic anemia. The cause of hypochromic microcytic anemia is typically decreased iron reserves in the body. This may be due to decreased dietary iron, poor absorption of iron from the gut, acute or chronic blood loss [42]. Iron absorption is primarily carried out by cells in the small intestine [43], explaining why it would be a top context for anemia. Obesity, which is an excessive accumulation of adipose, has also been molecularly linked to iron deficiency in a way that shows the conditions mutually affect one another [44]. Also of note, the primary affected tissue of pure red-cell aplasia, blood, was not identified in the top three contexts. However, patients with hepatitis, a symptom of liver inflammation, sometimes develop pure red-cell aplasia [45, 46]. CONE did manage to highlight cryptic associations between the liver and pure red-cell aplasia. Together, these results imply that CONE can help uncover subtle disease-tissue relationships. Thus, CONE contextualized embeddings can not only achieve good prediction performance, but the top performing contexts typically show biological relevance.

**Table 1.**
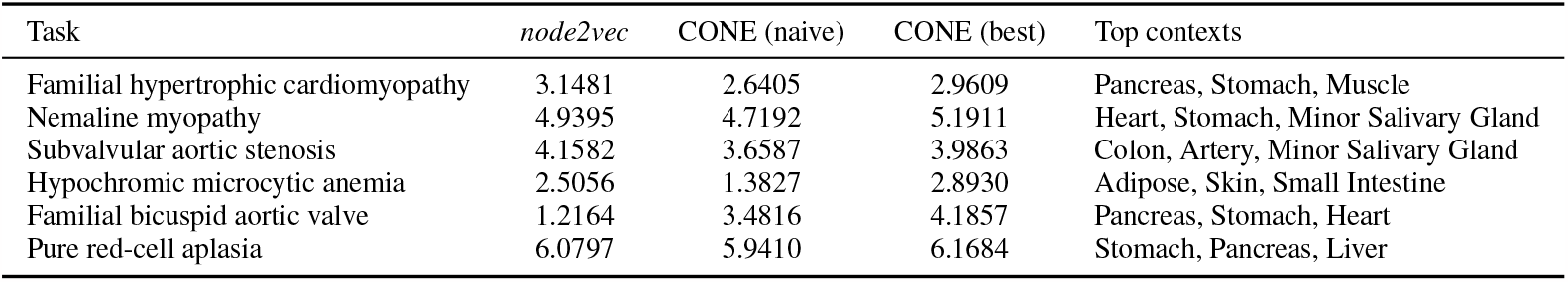
Top performing contexts for selected diseases in the DisGeNET benchmark. Performance reported as test APOP scores averaged across five random splits. The top contexts are sorted descendingly from left two right. For example, the Heart context achieved the highest score for Nemaline myopathy.

**Table 2.**
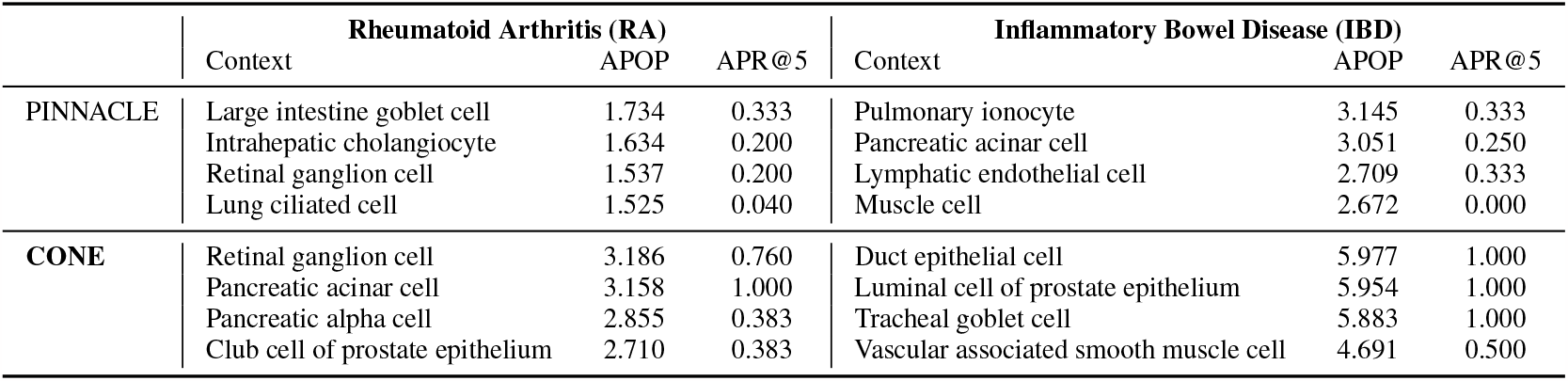
Top four performing contexts for predicting RA and IBD therapeutic targets. The first block of rows show the top four cell types with highest APOP socres of predicting RA and IBD using PINNACLE embeddings. Similarly, the second block of rows show those for the CONE embeddings.

|  | Rheumatoid Arthritis (RA) |  |  | Inflammatory Bowel Disease (IBD) |  |  |
| --- | --- | --- | --- | --- | --- | --- |
|  | Context | APOP | APR@5 | Context | APOP | APR@5 |
| PINNACLE | Large intestine goblet cell | 1.734 | 0.333 | Pulmonary ionocyte | 3.145 | 0.333 |
|  | Intrahepatic cholangiocyte | 1.634 | 0.200 | Pancreatic acinar cell | 3.051 | 0.250 |
|  | Retinal ganglion cell | 1.537 | 0.200 | Lymphatic endothelial cell | 2.709 | 0.333 |
|  | Lung ciliated cell | 1.525 | 0.040 | Muscle cell | 2.672 | 0.000 |
| CONE | Retinal ganglion cell | 3.186 | 0.760 | Duct epithelial cell | 5.977 | 1.000 |
|  | Pancreatic acinar cell | 3.158 | 1.000 | Luminal cell of prostate epithelium | 5.954 | 1.000 |
|  | Pancreatic alpha cell | 2.855 | 0.383 | Tracheal goblet cell | 5.883 | 1.000 |
|  | Club cell of prostate epithelium | 2.710 | 0.383 | Vascular associated smooth muscle cell | 4.691 | 0.500 |

#### RQ3. Can we further enhance therapeutic target prediction by encoding better contextual network information?

Inducing biological context information has been recently shown to be beneficial for predicting therapeutic targets in complex diseases such as Rheumatoid arthritis (RA) and Inflammatory Bowel Disease (IBD) [21]. Therapeutic target prediction for a particular disease is a binary classification problem that aims to predict whether a particular human gene can be used as a point of intervention for treating that disease. Compared to CONE, PINNACLE [21] takes an alternative approach by constructing a heterogeneous network that introduces different biological contexts as a type of node in the heterogeneous biomedical graph. Furthermore, the PINNACLE model learns individual sets of parameters for each biological context, contrasting with our unified model that shares the same set of parameters across all biological contexts. We hypothesize that our approach of tying weights induces more regularity from the underlying graph and, as a result, produces better contextualized embeddings for predicting therapeutic targets.

To test this, we first use the cell-type specific genes processed by PINNACLE to generate cell-type specific CONE embeddings. We then follow the original evaluation and measure different contextualized embeddings’ performance using APR@5. We note that the PINNACLE context-specific embeddings only contain embeddings of genes within the context. Conversely, CONE context-specific embeddings are genome-wide, meaning that embeddings in any context contain the same number of genes, spanning the whole network. Thus, to unify the setting between PINNACLE and CONE, we subset each context-specific CONE embedding to the corresponding context genes.

As shown in Figure 3, our CONE embeddings achieve significantly better performance than the PINNACLE embeddings when used in an unsupervised embedding setting, in which the training of the embedding does not involve node label information for the downstream prediction tasks. Furthermore, the highlighted immune cell contexts show that CONE better prioritizes the relevant cell context of IBD and RA, both of which are autoimmune diseases resulting from the malfunction of immune cells. Upon top-performing cell contexts in RA, CONE achieves better performance than PINNACLE. Specifically, CONE reveals contexts that are biologically related to RA. For example, pancreatic acinar cells (rank of APOP=2, rank of APR@5=1) secrete digestive enzymes that are involved in the digestion process within the small intestine. The early activation of these digestive enzymes before they reach the duodenum can trigger the onset of acute pancreatitis [47]. On the other hand, acute pancreatitis is highly associated with RA. Clinical studies have shown that RA patients were 2.51 times more likely to develop acute pancreatitis [48]. CONE is also able to reveal hidden associations based on cell-type-specific networks. Similarly, CONE performs better than PINNACLE in identifying top relevant cell contexts related to IBD. CONE picked up duct epithelial cells (rank of APOP=1, rank of APR@5=1) as the top cell context. These cells are integral to the intestinal barrier, serving as the first line of defense against invading microorganisms [49]. However, in IBD patients, the proper function of the intestinal barrier is frequently compromised to varying degrees [50]. Overall, these examples demonstrate the superior power of CONE in predicting therapeutic targets using biologically relevant cell contexts.

**Figure 3:**
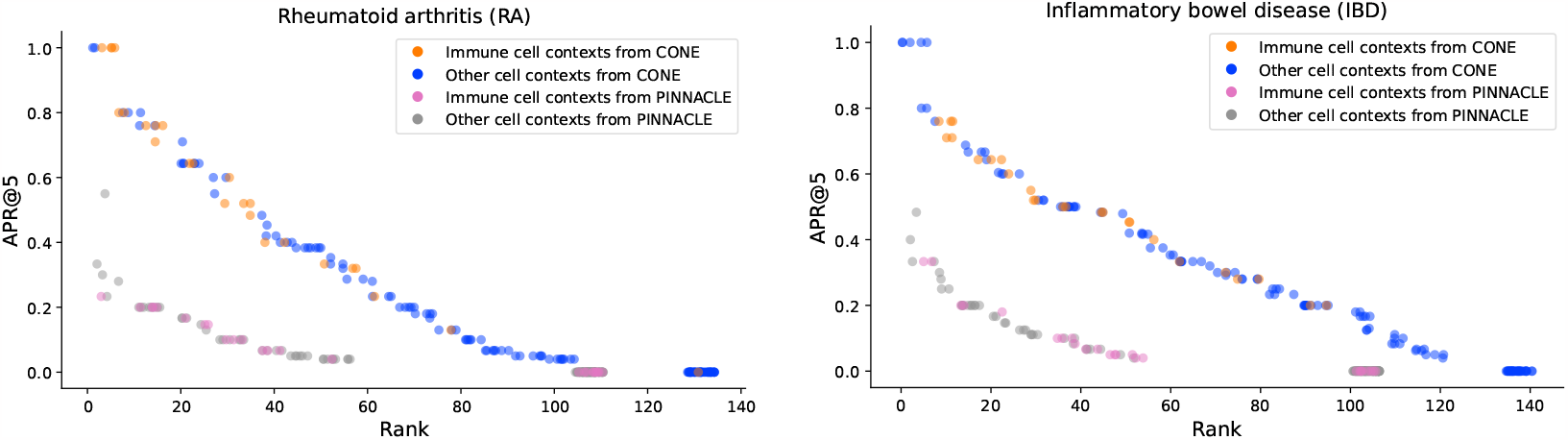
Therapeutic area predictions for RA and IBD. Each point represent the APR@5 score achieved when using a particular cell-type-specific embedding to predict the RA or IBD therapeutic targets. Immune cell contexts are highlighted in orange and pink for CONE and PINNACLE.

#### RQ4. Can CONE leverage biological contexts other than tissue or cell-type?

Since CONE takes contextual information in the form of node sets, it can be extended to biological contexts outside of traditionally used tissue and cell type contexts [20, 21]. Here, we explore the extendability of our CONE approach to different biological contexts in two other ways. First, we reevaluate the RA and IBD prediction performance using CONE trained on different disease contexts defined by differentially expressed genes obtained from CREEDS [51]. We find that the top-ranked contexts for both RA and IBD are indeed highly relevant disorders for both diseases (Figure 4). For example, psoriasis is one of the top disease contexts related to RA (rank of APOP=3). A clear connection between these two conditions is psoriatic arthritis, a form of arthritis with a skin rash, which is a common symptom in psoriasis [52]. This indicates that there are similar genetic programs shared by both diseases [53], which is revealed by CONE using disease context networks. Furthermore, CONE also reveals several connections between RA and some seemingly unrelated diseases, such as heart failure (rank of APOP=115). Notably, a recent study has confirmed that RA patients have a two-fold higher risk of heart failure mortality than those without RA [54].

**Figure 4:**
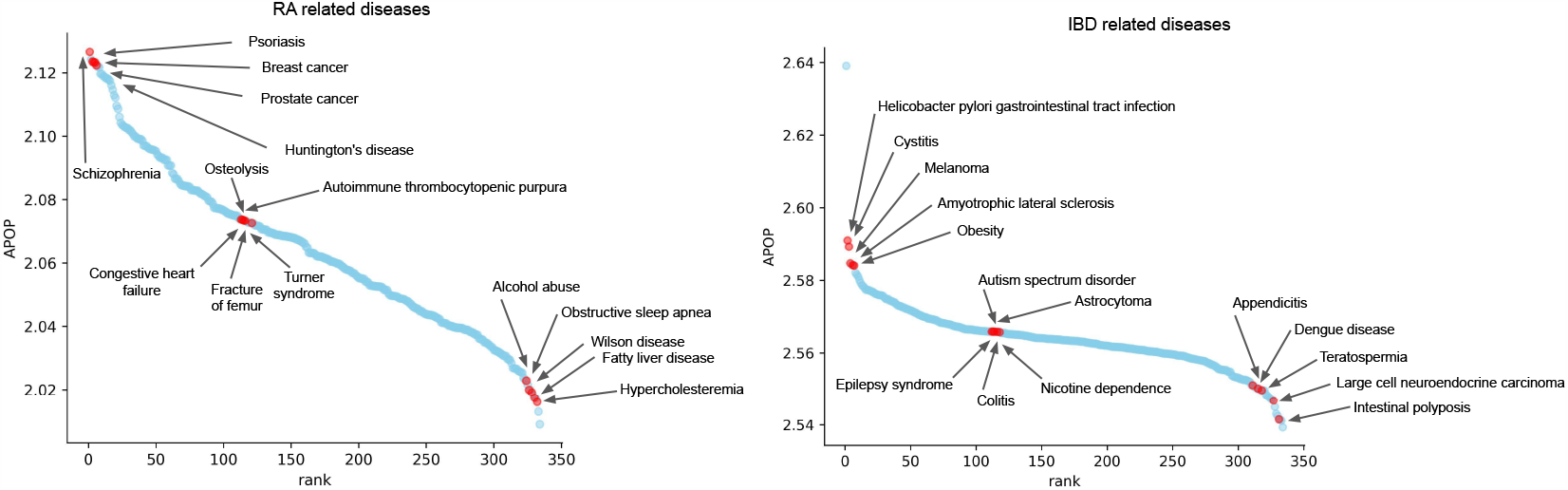
Sorted disease contexts for predict the therapeutic area predictions for RA and IBD.

Similarly, CONE unveils meaningful relationships between IBD and other disease contexts. Cystitis (rank of APOP=3) is one of the top disease contexts identified by CONE. Clinical studies have shown that cystitis, an inflammation of the bladder, leads to a significant increase in the risk of IBD [55–57]. For example, individuals with interstitial cystitis, a condition involving an inflamed or irritated bladder wall, are 100 times more likely to have IBD [57]. CONE also finds subtle relationships between IBD and neurological complications exemplified by epilepsy syndrome (rank of APOP=114) and autism spectrum disorder (rank of APOP=112) in the top list [58]. Neurological complications affect 0.25% to 47.50% IBD patients [59]. These IBD-related neurological complications are associated with neuroinflammation or increased risk of blood clots in brain veins [60]. Some diseases may have a protective effect on other diseases. CONE identified such protective relationships between *Helicobacter pylori* gastrointestinal tract infection (rank of APOP=2) and IBD, since *H. pylori* infection helps protect against IBD by inducing systematic immune tolerance and suppressing inflammatory responses [61]. Overall, these examples further confirm that CONE can leverage disease context to reveal both apparent and cryptic associations between complex diseases.

Finally, we reperform the DisGeNET benchmark using a diverse construction of context specific gene sets, spanning from tissues, cell types, to diseases, and retrieved from various databases, including CellMarker2.0 [62], Human Protein Atlas [63], and the TISSUES database [64]. We observe that CONE performs similarly under all collections of contexts tested (Figure B.3). Together with the fact that CONE performs competitively against baselines (***RQ1***), and that it captures biologically meaningful contextual information (***RQ2***), we believe that CONE is a versatile and effective approach to scalably generate biological network embeddings conditioned on specific biological contexts.

#### RQ5. Cane CONE transfer to unseen contexts?

The MLP context encoder used by CONE provides possibility to generate embeddings for contexts that are not observed during training. This can be done by feeding the similarity profile of the new query context against all training contexts to the MLP encoder. To demonstrate the effectiveness of transfering to unseen context, we retrain the CONE GTEx tissue-specific embeddings, but leaving out the Heart context during training. We observe that holding out Heart context does not affect the disease gene classification performance significantly, with *>* 0.8 coefficient of correlation against the original performance. Furthermore, we compile two lists of diseases for which the Heart context achieved top five performances, for the riginal and held-out Heart versions of CONE. We found a significant overlap between the two lists (Hypergeometric p-val *<* 0.05), with seven common diseases (Table B.2). One notable example is familial bicuspid aortic valve, a known common congenital heart defect [65]. These results highlight the effective transferability of CONE to unseen contexts, with similar embedding quality as if the contexts were seen during training.

## 6 Conclusion

We proposed CONE, a flexible and scalable framework that can inject arbitrary contextual information into gene interaction networks. Our study underscores the efficacy of the CONE method in enhancing the prioritization of genes within the gene interaction network. CONE consistently demonstrated superior performance in various biomedical tasks than baseline methods. Crucially, the introduction of context-specific embeddings, especially when positive genes for a disease were limited, resulted in significant performance gains. Moreover, the contexts identified by CONE were found to be biologically relevant, suggesting that the method not only boosts prediction accuracy but also provides biologically meaningful insights. This ability to integrate diverse biological contexts, from tissues and cell types to diseases, positions CONE as a versatile tool that can be used to uncover both explicit and cryptic relationships within biomedical datasets.

### Limitations and future directions

Constructing context-specific subnetworks solely based on the subgraph induced by context-specific genes is a reasonable but overly simplistic assumption. In reality, context-specific gene interactions are complicated and encompass diverse mechanisms, ranging from interactions mediated by non-coding RNAs to the influence of epigenetic modifications and signaling pathways. Consequently, a promising advancement would involve the carefully constructed context-specific gene interaction networks that account for these intricate nuances, such as HumanBase [66], HIPPIE [67], IID [68], and many others [69–71].

## A Scalability experiment

We empirically demonstrate the scalability of CONE against three other related methods, including GAT, BIONIC, and PINNACLE. All these methods use the GAT module as the main encoding component. CONE and GAT both employ a single GAT module, while BIONIC and PINNACLE use individual GAT modules for different contexts.

### Setup

We consider two types of scaling experiments, *number of contexts* and *context node percentages*. For *number of contexts*, we fix the number of context-specific nodes to be about 5% of the total number of nodes and vary the number of contexts from 10 to 1000. Meanwhile, for *context node percentages*, we fix the number of contexts to be 100, and vary the context node percentages from 1% to 50%. For both sets of experiments, we use a synthetic network with 10, 000 nodes generated using the Barabási–Albert model [72]. The network is generated so that the density is approximately 0.01. We report the peak CUDA memory usage (in bytes) of the forward pass for the model using the `torch.cuda.max_memory_allocated()` function.

### Implementation details

All experiments are conducted on compute nodes with 5 CPUs, 45GB memory, and a Tesla v100 GPU (32GB). We uniformly set the following hyperparameters across models: 128 dimensions and one layer. Furthermore, BIONIC and PINNACLE require subgraph batched training. We set the batch size to 2048 for both models. On the other hand, CONE and GAT employ full batched computation.

### Results

The empirical scalability results are shown in Figure A.1. We first highlight that CONE shows great scalability, with very minimal overhead as more contexts are introduced. Remarkably, CONE’s memory consumption is quite comparable to the plain GAT model that does not take context into account. This aligns well with our complexity analysis (Section 4.4).

Conversely, BIONIC and PINNACLE react drastically to the number of contexts, with BIONIC running out of memory beyond 500 contexts. Furthermore, PINNACLE demonstrates a severe scalability issue with the context nodes percentage. In other words, PINNACLE memory consumption drastically increases as the context subgraphs increase. These results showcase the scalability advantage of CONE using a single shared GAT module to decode embeddings for all contexts.

**Figure A.1:**
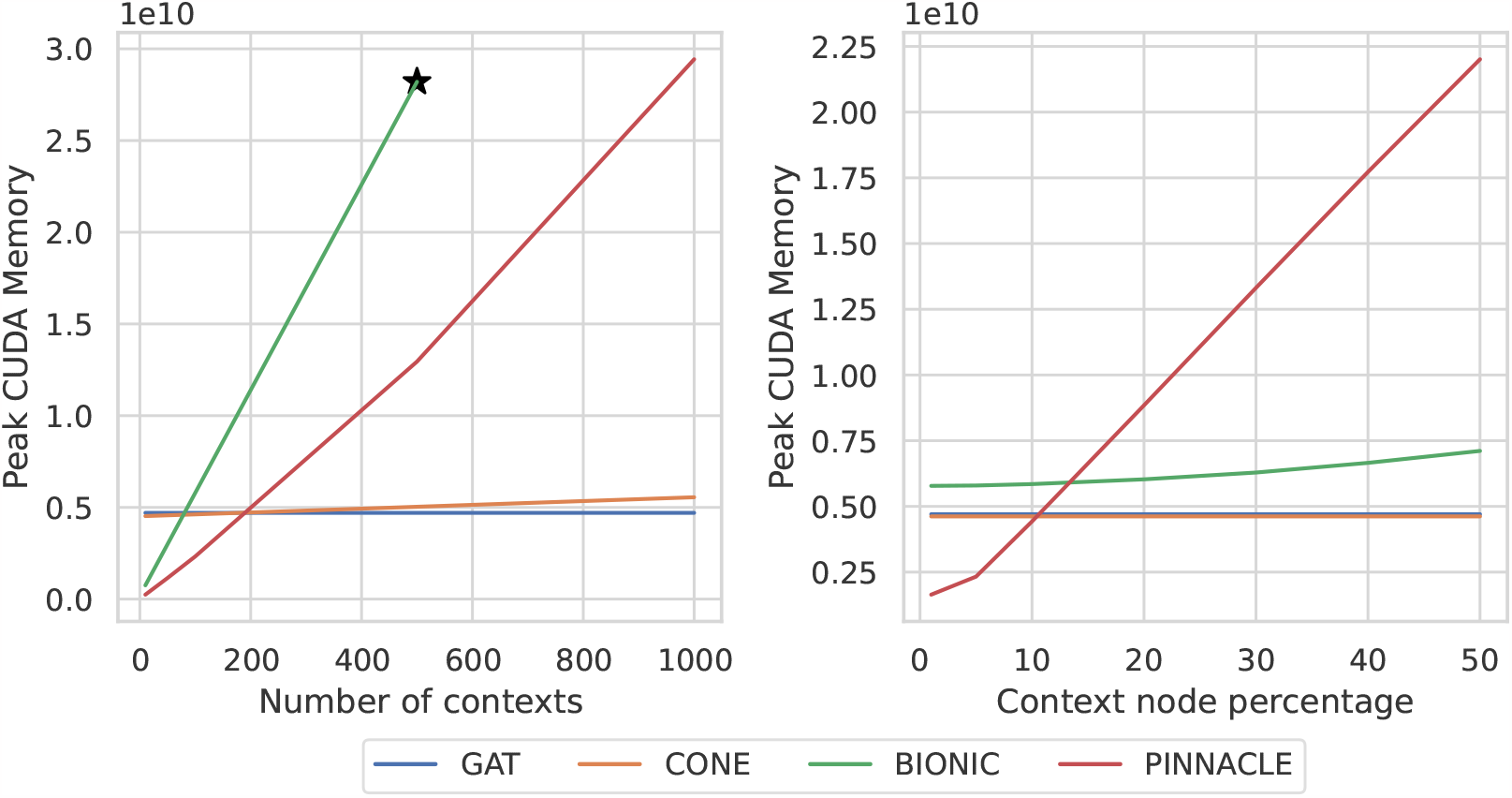
Models scalability across different contextualization settings. Star indicates the point beyond which the model will run out of memory.

## B Additional results

### B.1 Effects of PCA dimensionality reduction

The final CONE context-specific embeddings are obtained by first concatenating the context-naive with the context-specific embeddings, and then applying PCA to project the dimension by half. Combining the context-naive and context-specific embeddings gives the final embedding a more comprehensive view of both the global and local (context-specific) semantics. Dimensionality reduction is applied so that the results for the final context-specific embeddings can be fairly compared to the context-naive embeddings. PCA is a common dimensionality reduction technique due to its simplicity and has been used in previous studies to combine multiple views of the embeddings, such as Walklets [73].

One question remaining is how does the performance change before and after applying PCA. Here, we compare the performance between the fully concatenated and PCA-reduced versions of CONE following the DisGeNET benchmarking setting in ***RQ1***. We observe little performance difference between the two versions of CONE (Figure B.1). Thus, we set the final CONE to use PCA, as it provides a fairer setting in terms of the number of dimensions while having the same performance.

**Figure B.1:**
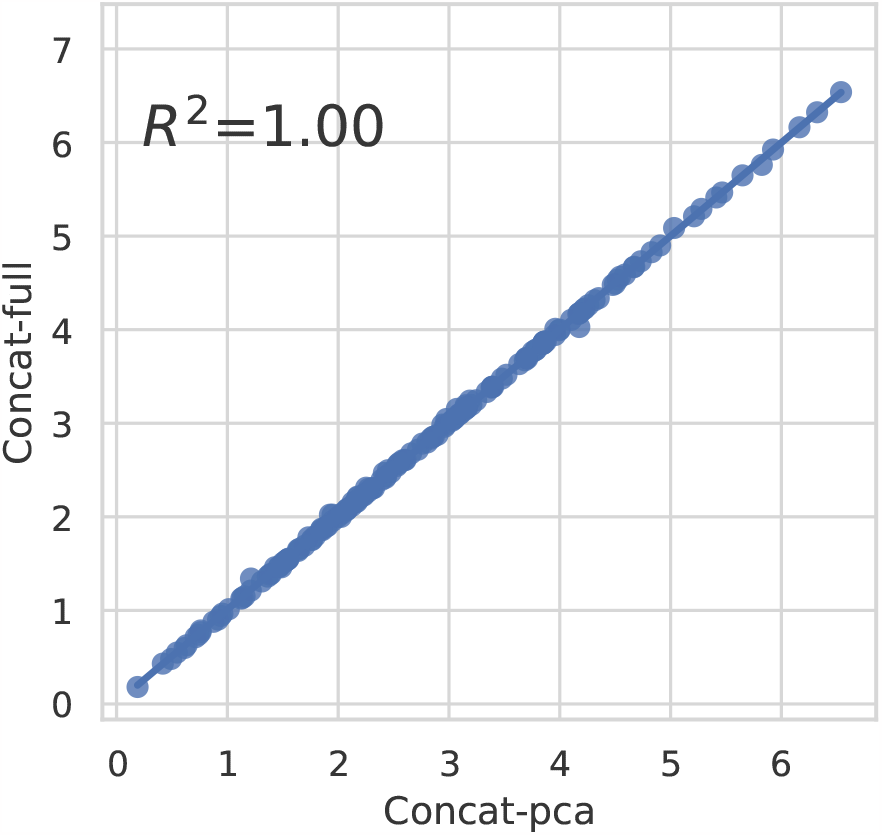
CONE dimensionality reduction effect comparison. Performance comparison between PCA-reduced CONE (x-axis) and not reduced CONE (y-axis) in terms of testing APOP on the DisGeNET disease gene classification benchmark.

### B.2 Ablation studies

In the followings, we investigate the effectiveness of our main design choices of CONE, including the context similarity measure and the MLP context encoder. We follow the same experimental settings in ***RQ1*** using the DisGeNET gene classification benchmark.

#### Context similarity measures

Besides the default Jaccard similarity measure, we consider three other similarity measures, including the cosine similarity, radial basis function, and Spearman correlation coefficient. Table B.1 shows that the choice of similarity has marginal effects on the performance, with the default Jaccard similarity consistently achieving better or equivalent performance against other choices of similarities. Figure B.2 further indicate that there is no significant performance differences across the choice of similarity measures according to the paired Wilcoxon test.

#### Context encoder

A trivial way to encode context embedding is one-hot encoding, which is equivalent to directly learning the embedding for each contexts. We call this approach *Embedding*. We observe that *Embedding* achieves the lowest average performance across all groups of tasks (Table B.1). In the case of disease task group [23, 42), the *Embedding* performance is significantly worse than that of the default CONE (Figure B.2, Wilcoxon p-value *<* 0.05).

**Table B.1:**
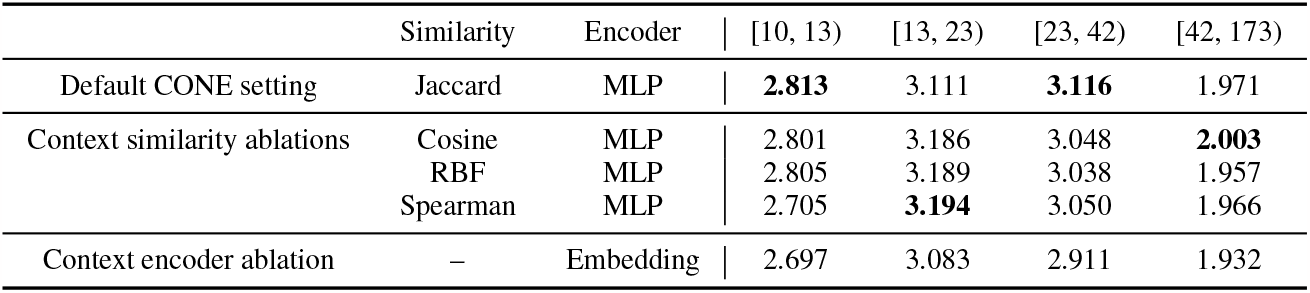
Ablation study of context similarity and context encoding strategies using the DisGeNET benchmark. Results are reported as APOP scores averaged across tasks within a group based on the number of positive examples.

**Figure B.2:**
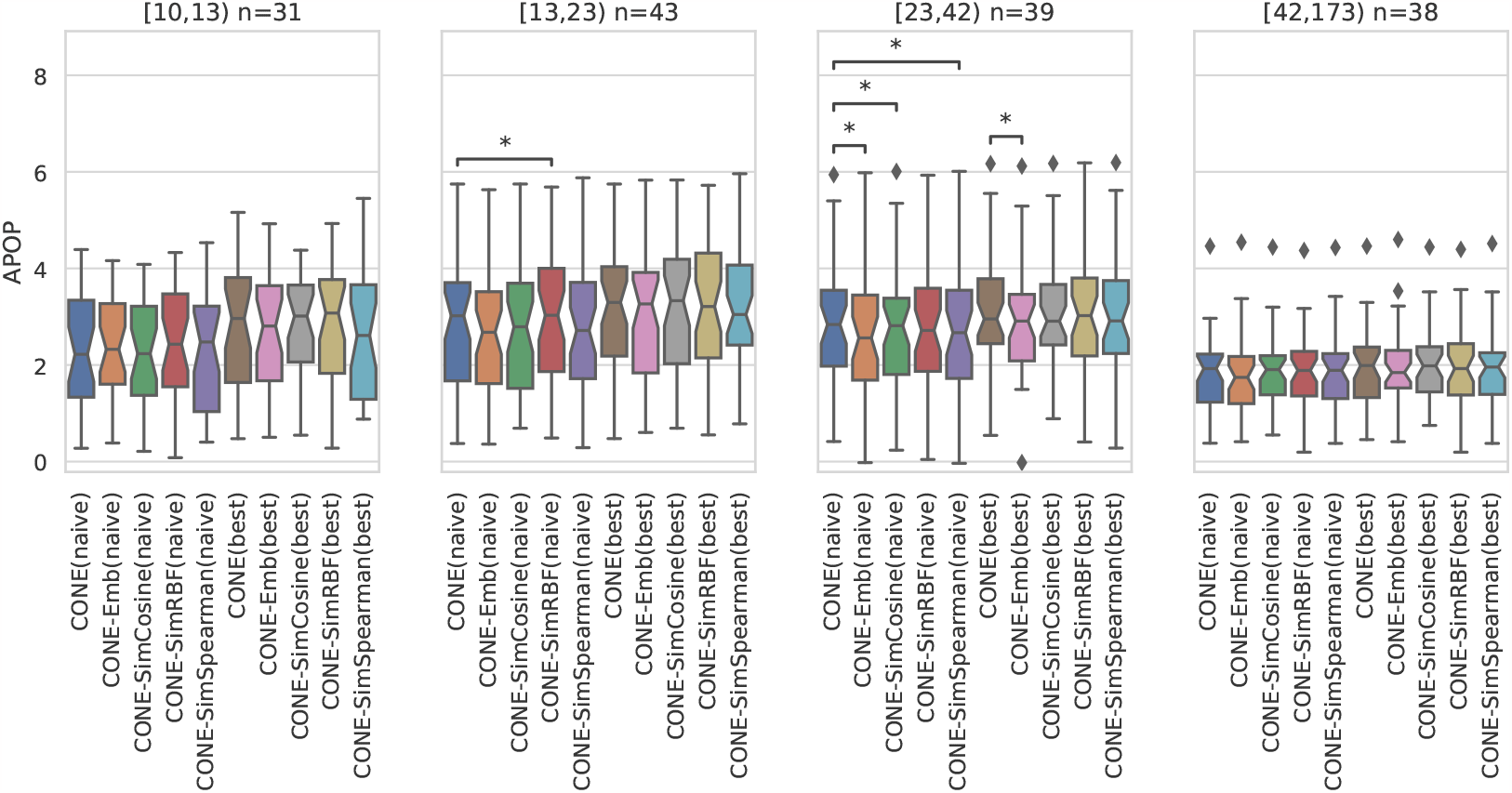
Performance for different context similarity and context encoding strategies. Each point in the box plot corresponds to the prediction test performance of a disease gene classification task from the DisGeNET benchmark, averaged across five random splits. Different panels show groups of diseases with different number of positive genes. * indicate that the performances between the two methods are significantly different (Wilcoxon p-value *<* 0.05).

**Figure B.3:**
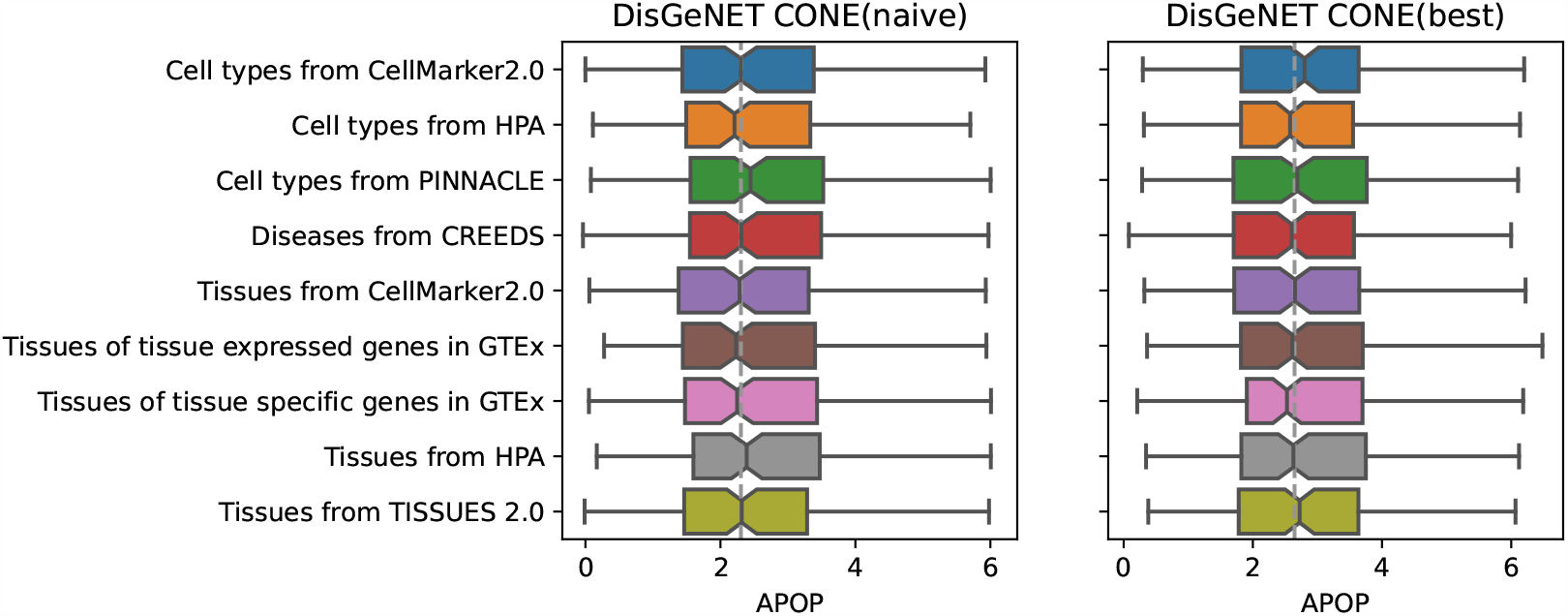
Performance of CONE on the DisGeNET benchmark across contexts collections.

**Table B.2:**
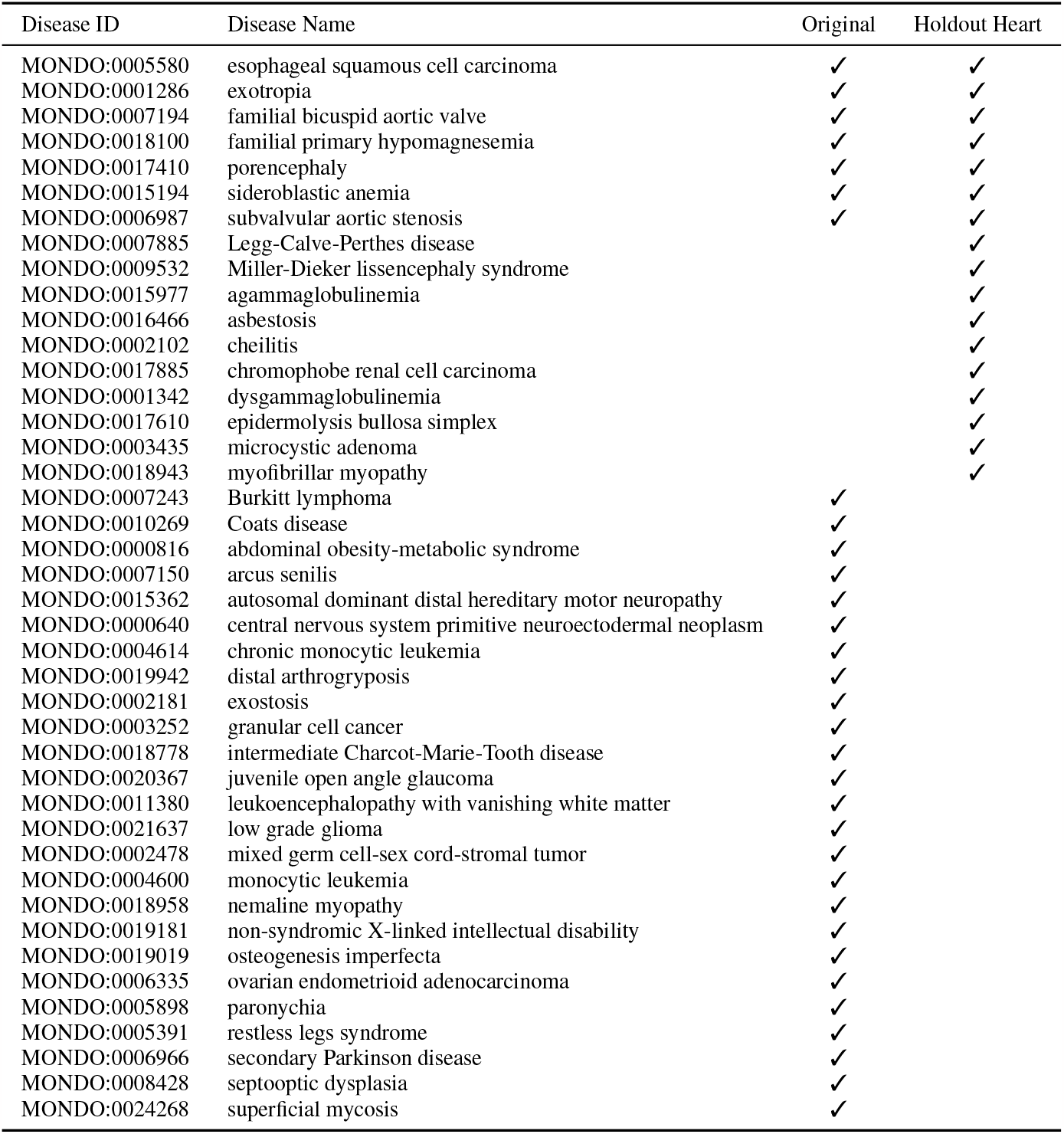
List of diseases for which the Heart context appears to be the top five performing contexts in terms of test APOP scores. Last two columns indicate whether the disease shows up as top five context for the original CONE or the one trained without Heart context.

## C Additional information

### C.1 Dataset information

#### Network

We obtain the raw PINPPI network from the PINNACLE paper^5^, which contains 15, 461 nodes and 207, 641 edges. We then convert the node IDs from gene symbol to Entrez ID [74] using the MyGeneInfo query service [75]. We only perserve genes that have exact one-to-one mapping from gene symbol to Entrez ID. After the above conversion, the final processed Entrez based PINPPI network contains 15, 229 nodes and 206, 835 edges.

**Table C.1:** Gene set statistics. First three gene set collections are used as prediction tasks, and the remaining three gene set collections are used as contexts.

|  | Collection | Type | # gene sets | Min. | Lower quart. | Med. | Upper quart. | Max. |
| --- | --- | --- | --- | --- | --- | --- | --- | --- |
| Tasks | DisGeNET [36] | Disease | 167 | 10 | 13 | 21 | 39 | 189 |
|  | IBD [21, 76] | Therapeutic Target | 1 |  |  | 151 |  |  |
|  | RA [21, 76] | Therapeutic Target | 1 |  |  | 113 |  |  |
| Contexts | GTEX [40] | Tissue | 30 | 430 | 1122.5 | 1593.5 | 3091 | 8604 |
|  | Tabula Sapiens [21, 22] | Cell type | 156 | 1002 | 2012 | 2751 | 3051 | 3425 |
|  | CREEDS [51] | Disease | 333 | 204 | 535 | 566 | 998 | 4452 |

<sup>5</sup>https://figshare.com/articles/software/PINNACLE/22708126

**Table C.2:** Hyperparameter search grid. Final settings are **bolded**.

|  | Hyperparameter | Search grid |
| --- | --- | --- |
| Architecture | Number of layers | [1, **2**, 3] |
|  | Number of heads | [1, 2, **3**, 4, 5, 6] |
| Optimization | Learning rate | [0.01, 0.005, **0.001**, 0.0005, 0.0001] |
|  | Weight decay | [1, 0.5, **0.1**, 0.05, 0.001, 0.0005, 0.0001] |
| Random walk | Walk length | [40, 80, **120**, 160, 200] |
|  | Walks per node | [1, 5, **10**, 15, 20] |
|  | Window size | [5, **10**, 15, 20, 25] |

### C.2 Hyperparameter tuning

We optimized CONE’s hyperparameters using our main DisGeNET benchmark (***RQ1***) based on the averaged validation APOP scores of the context-naive CONE embeddings. The hyperparameter grid is shown in Table C.2, where the final hyperparameter settings are bolded.

### Footnotes

3 https://github.com/krishnanlab/cone

4 https://figshare.com/articles/software/PINNACLE/22708126

5 https://figshare.com/articles/software/PINNACLE/22708126

## References

[1] Albert-László Barabási, Natali Gulbahce, and Joseph Loscalzo. Network medicine: a network-based approach to human disease. Nature reviews genetics, 12(1):56–68, 2011. 1

[2] Marc Vidal, Michael E Cusick, and Albert-László Barabási. Interactome networks and human disease. Cell, 144(6):986–998, 2011. 1

[3] Renming Liu, Christopher A Mancuso, Anna Yannakopoulos, Kayla A Johnson, and Arjun Krishnan. Supervised learning is an accurate method for network-based gene classification. Bioinformatics, 36(11):3457–3465, 2020. 1, 5, 6

[4] Arjun Krishnan, Ran Zhang, Victoria Yao, Chandra L Theesfeld, Aaron K Wong, Alicja Tadych, Natalia Volfovsky, Alan Packer, Alex Lash, and Olga G Troyanskaya. Genome-wide prediction and functional characterization of the genetic basis of autism spectrum disorder. Nature neuroscience, 19(11):1454–1462, 2016. 1

[5] Victoria Yao, Rachel Kaletsky, William Keyes, Danielle E Mor, Aaron K Wong, Salman Sohrabi, Coleen T Murphy, and Olga G Troyanskaya. An integrative tissue-network approach to identify and test human disease genes. Nature biotechnology, 36(11):1091–1099, 2018. 1

[6] Xiang Yue, Zhen Wang, Jingong Huang, Srinivasan Parthasarathy, Soheil Moosavinasab, Yungui Huang, Simon M Lin, Wen Zhang, Ping Zhang, and Huan Sun. Graph embedding on biomedical networks: methods, applications and evaluations. Bioinformatics, 36(4):1241–1251, 2020. 1

[7] Dongsheng Duan, Nathalie Goemans, Shin’ichi Takeda, Eugenio Mercuri, and Annemieke Aartsma-Rus. Duchenne muscular dystrophy. Nature Reviews Disease Primers, 7(1):13, 2021. 1

[8] Mary Lauren Benton, Abin Abraham, Abigail L LaBella, Patrick Abbot, Antonis Rokas, and John A Capra. The influence of evolutionary history on human health and disease. Nature Reviews Genetics, 22(5):269–283, 2021. 1

[9] Idan Hekselman and Esti Yeger-Lotem. Mechanisms of tissue and cell-type specificity in heritable traits and diseases. Nature Reviews Genetics, 21(3):137–150, 2020. 2

[10] Myat Noe Han, David I Finkelstein, Rachel M McQuade, and Shanti Diwakarla. Gastrointestinal dysfunction in parkinson’s disease: current and potential therapeutics. Journal of Personalized Medicine, 12(2):144, 2022. 2

[11] Sarvenaz Choobdar, Mehmet E Ahsen, Jake Crawford, Mattia Tomasoni, Tao Fang, David Lamparter, Junyuan Lin, Benjamin Hescott, Xiaozhe Hu, Johnathan Mercer, et al. Assessment of network module identification across complex diseases. Nature methods, 16(9):843–852, 2019. 2

[12] Rose Oughtred, Jennifer Rust, Christie Chang, Bobby-Joe Breitkreutz, Chris Stark, Andrew Willems, Lorrie Boucher, Genie Leung, Nadine Kolas, Frederick Zhang, et al. The biogrid database: A comprehensive biomedical resource of curated protein, genetic, and chemical interactions. Protein Science, 30(1):187–200, 2021. 2

[13] Damian Szklarczyk, Andrea Franceschini, Stefan Wyder, Kristoffer Forslund, Davide Heller, Jaime Huerta-Cepas, Milan Simonovic, Alexander Roth, Alberto Santos, Kalliopi P Tsafou, et al. String v10: protein–protein interaction networks, integrated over the tree of life. Nucleic acids research, 43(D1):D447–D452, 2015.

[14] Chan Yeong Kim, Seungbyn Baek, Junha Cha, Sunmo Yang, Eiru Kim, Edward M Marcotte, Traver Hart, and Insuk Lee. Humannet v3: an improved database of human gene networks for disease research. Nucleic acids research, 50(D1):D632–D639, 2022. 2

[15] Vladimir Gligorijević, Meet Barot, and Richard Bonneau. deepnf: deep network fusion for protein function prediction. Bioinformatics, 34(22):3873–3881, 2018. 2

[16] Duncan T Forster, Sheena C Li, Yoko Yashiroda, Mami Yoshimura, Zhijian Li, Luis Alberto Vega Isuhuaylas, Kaori Itto-Nakama, Daisuke Yamanaka, Yoshikazu Ohya, Hiroyuki Osada, et al. Bionic: biological network integration using convolutions. Nature Methods, 19 (10):1250–1261, 2022. 2, 5

[17] Patrick Danaher, Pei Wang, and Daniela M Witten. The joint graphical lasso for inverse covariance estimation across multiple classes. Journal of the Royal Statistical Society Series B: Statistical Methodology, 76(2):373–397, 2014.

[18] Hyunghoon Cho, Bonnie Berger, and Jian Peng. Compact integration of multi-network topology for functional analysis of genes. Cell systems, 3(6):540–548, 2016.

[19] Adelaide Woicik, Mingxin Zhang, Hanwen Xu, Sara Mostafavi, and Sheng Wang. Gemini: memory-efficient integration of hundreds of gene networks with high-order pooling. Bioinformatics, 39:i504–i512, 2023. 2, 5

[20] Marinka Zitnik and Jure Leskovec. Predicting multicellular function through multi-layer tissue networks. Bioinformatics, 33(14):i190–i198, 2017. 2, 8

[21] Michelle M Li, Yepeng Huang, Marissa Sumathipala, Man Qing Liang, Alberto Valdeolivas, Ashwin N Ananthakrishnan, Katherine Liao, Daniel Marbach, and Marinka Zitnik. Contextualizing protein representations using deep learning on protein networks and single-cell data. bioRxiv,pages 2023–07, 2023. 2, 5, 7, 8, 18

[22] Tabula Sapiens Consortium*, Robert C Jones, Jim Karkanias, Mark A Krasnow, Angela Oliveira Pisco, Stephen R Quake, Julia Salzman, Nir Yosef, Bryan Bulthaup, Phillip Brown, et al. The tabula sapiens: A multiple-organ, single-cell transcriptomic atlas of humans. Science, 376 (6594):eabl4896, 2022. 2, 18

[23] Sami Abu-El-Haija, Bryan Perozzi, Rami Al-Rfou, and Alexander A Alemi. Watch your step: Learning node embeddings via graph attention. Advances in neural information processing systems, 31, 2018. 2

[24] Aditya Grover and Jure Leskovec. node2vec: Scalable feature learning for networks. In Proceedings of the 22nd ACM SIGKDD international conference on Knowledge discovery and data mining, pages 855–864, 2016. 3

[25] Michael Gutmann and Aapo Hyvärinen. Noise-contrastive estimation: A new estimation principle for unnormalized statistical models. In Proceedings of the thirteenth international conference on artificial intelligence and statistics, pages 297–304. JMLR Workshop and Conference Proceedings, 2010. 3

[26] Zonghan Wu, Shirui Pan, Fengwen Chen, Guodong Long, Chengqi Zhang, and S Yu Philip. A comprehensive survey on graph neural networks. IEEE transactions on neural networks and learning systems, 32(1):4–24, 2020. 3

[27] Jie Zhou, Ganqu Cui, Shengding Hu, Zhengyan Zhang, Cheng Yang, Zhiyuan Liu, Lifeng Wang, Changcheng Li, and Maosong Sun. Graph neural networks: A review of methods and applications. AI open, 1:57–81, 2020. 3

[28] Petar Veličković, Guillem Cucurull, Arantxa Casanova, Adriana Romero, Pietro Liò, and Yoshua Bengio. Graph attention networks. In International Conference on Learning Representations, 2018. 3, 5

[29] Shaked Brody, Uri Alon, and Eran Yahav. How attentive are graph attention networks? In International Conference on Learning Representations, 2021. 3, 5

[30] Ashish Vaswani, Noam Shazeer, Niki Parmar, Jakob Uszkoreit, Llion Jones, Aidan N Gomez, Lukasz Kaiser, and Illia Polosukhin. Attention is all you need. Advances in neural information processing systems, 30, 2017. 3

[31] Robin Rombach, Andreas Blattmann, Dominik Lorenz, Patrick Esser, and Björn Ommer. Highresolution image synthesis with latent diffusion models. In Proceedings of the IEEE/CVF conference on computer vision and pattern recognition, pages 10684–10695, 2022. 4

[32] Ilya Loshchilov and Frank Hutter. Decoupled weight decay regularization. In International Conference on Learning Representations, 2018. 4

[33] Chris Stark, Bobby-Joe Breitkreutz, Teresa Reguly, Lorrie Boucher, Ashton Breitkreutz, and Mike Tyers. Biogrid: a general repository for interaction datasets. Nucleic acids research, 34 (suppl_1):D535–D539, 2006. 5

[34] Jörg Menche, Amitabh Sharma, Maksim Kitsak, Susan Dina Ghiassian, Marc Vidal, Joseph Loscalzo, and Albert-László Barabási. Uncovering disease-disease relationships through the incomplete interactome. Science, 347(6224):1257601, 2015. 5

[35] Katja Luck, Dae-Kyum Kim, Luke Lambourne, Kerstin Spirohn, Bridget E Begg, Wenting Bian, Ruth Brignall, Tiziana Cafarelli, Francisco J Campos-Laborie, Benoit Charloteaux, et al. A reference map of the human binary protein interactome. Nature, 580(7803):402–408, 2020. 5

[36] Janet Piñero, Àlex Bravo, Núria Queralt-Rosinach, Alba Gutiérrez-Sacristán, Jordi Deu-Pons, Emilio Centeno, Javier García-García, Ferran Sanz, and Laura I Furlong. Disgenet: a comprehensive platform integrating information on human disease-associated genes and variants. Nucleic acids research, page gkw943, 2016. 5, 18

[37] Sezin Kircali Ata, Min Wu, Yuan Fang, L. Ou-Yang, Chee Keong Kwoh, and Xiao-Li Li. Recent advances in network-based methods for disease gene prediction. Briefings in bioinformatics, 22 (4):bbaa303, 2021. 5

[38] Thomas N Kipf and Max Welling. Variational graph auto-encoders. NIPS Workshop on Bayesian Deep Learning, 2016. 5

[39] Cole Trapnell. Defining cell types and states with single-cell genomics. Genome research, 25 (10):1491–1498, 2015. 5

[40] John Lonsdale, Jeffrey Thomas, Mike Salvatore, Rebecca Phillips, Edmund Lo, Saboor Shad, Richard Hasz, Gary Walters, Fernando Garcia, Nancy Young, et al. The genotype-tissue expression (gtex) project. Nature genetics, 45(6):580–585, 2013. 5, 18

[41] Takaya Saito and Marc Rehmsmeier. The precision-recall plot is more informative than the roc plot when evaluating binary classifiers on imbalanced datasets. PloS one, 10(3):e0118432, 2015. 6

[42] Hammad S. Chaudhry and Madhukar Reddy Kasarla. Microcytic Hypochromic Anemia. StatPearls Publishing, Treasure Island (FL), 2022. URL http://europepmc.org/books/NBK470252. 7

[43] Elif Piskin, Danila Cianciosi, Sukru Gulec, Merve Tomas, and Esra Capanoglu. Iron absorption: factors, limitations, and improvement methods. ACS omega, 7(24):20441–20456, 2022. 7

[44] Elmar Aigner, Alexandra Feldman, and Christian Datz. Obesity as an emerging risk factor for iron deficiency. Nutrients, 6(9):3587–3600, 2014. 7

[45] Pyoung Suk Lim, In Hee Kim, Seong Hun Kim, Seung Ok Lee, and Sang Wook Kim. A case of severe acute hepatitis a complicated with pure red cell aplasia. The Korean Journal of Gastroenterology, 60(3):177–181, 2012. 7

[46] Akira Sato, Fumiaki Sano, Toshiya Ishii, Kayo Adachi, Ryujirou Negishi, Nobuyuki Matsumoto, and Chiaki Okuse. Pure red cell aplasia associated with autoimmune hepatitis successfully treated with cyclosporine a. Clinical Journal of Gastroenterology, 7:74–78, 2014. 7

[47] Po Sing Leung and Siu Po Ip. Pancreatic acinar cell: its role in acute pancreatitis. The international journal of biochemistry & cell biology, 38(7):1024–1030, 2006. 8

[48] Motasem Alkhayyat, Mohannad Abou Saleh, Mehnaj Kaur Grewal, Mohammad Abureesh, Emad Mansoor, C Roberto Simons-Linares, Abby Abelson, and Prabhleen Chahal. Pancreatic manifestations in rheumatoid arthritis: a national population-based study. Rheumatology, 60(5): 2366–2374, 2021. 8

[49] Giulia Roda, Alessandro Sartini, Elisabetta Zambon, Andrea Calafiore, Margherita Marocchi, Alessandra Caponi, Andrea Belluzzi, and Enrico Roda. Intestinal epithelial cells in inflammatory bowel diseases. World journal of gastroenterology: WJG, 16(34):4264, 2010. 8

[50] Lena Antoni, Sabine Nuding, Jan Wehkamp, and Eduard F Stange. Intestinal barrier in inflammatory bowel disease. World journal of gastroenterology: WJG, 20(5):1165, 2014. 8

[51] Zichen Wang, Caroline D Monteiro, Kathleen M Jagodnik, Nicolas F Fernandez, Gregory W Gundersen, Andrew D Rouillard, Sherry L Jenkins, Axel S Feldmann, Kevin S Hu, Michael G McDermott, et al. Extraction and analysis of signatures from the gene expression omnibus by the crowd. Nature communications, 7(1):12846, 2016. 8, 18

[52] Keiichi Yamanaka, Osamu Yamamoto, and Tetsuya Honda. Pathophysiology of psoriasis: A review. The Journal of dermatology, 48(6):722–731, 2021. 8

[53] Liz Parrish. Psoriasis: symptoms, treatments and its impact on quality of life. British Journal of Community Nursing, 17(11):524–528, 2012. 8

[54] Elizabeth Park, Jan Griffin, and Joan M Bathon. Myocardial dysfunction and heart failure in rheumatoid arthritis. Arthritis & Rheumatology, 74(2):184–199, 2022. 9

[55] SM Mahmudul Hasan and Baljinder S Salh. Emphysematous cystitis as a potential marker of severe crohn’s disease. BMC gastroenterology, 22(1):181, 2022. 9

[56] Doreen E Chung, Lesley K Carr, Linda Sugar, Michelle Hladunewich, and Leslie A Deane. Xanthogranulomatous cystitis associated with inflammatory bowel disease. Canadian Urological Association Journal, 4(4):E91, 2010.

[57] Madhu Alagiri, Sherman Chottiner, Vicki Ratner, Debra Slade, and Philip M Hanno. Interstitial cystitis: unexplained associations with other chronic disease and pain syndromes. Urology, 49 (5):52–57, 1997. 9

[58] Giovanni Casella, Gian Eugenio Tontini, Gabrio Bassotti, Luca Pastorelli, Vincenzo Villanacci, Luisa Spina, Vittorio Baldini, and Maurizio Vecchi. Neurological disorders and inflammatory bowel diseases. World Journal of Gastroenterology: WJG, 20(27):8764, 2014. 9

[59] José M Ferro and Miguel Oliveira Santos. Neurology of inflammatory bowel disease. Journal of the Neurological Sciences, 424:117426, 2021. 9

[60] Germán Morís. Inflammatory bowel disease: an increased risk factor for neurologic complications. World Journal of Gastroenterology: WJG, 20(5):1228, 2014. 9

[61] Yang Yu, Shengtao Zhu, Peng Li, Li Min, and Shutian Zhang. Helicobacter pylori infection and inflammatory bowel disease: a crosstalk between upper and lower digestive tract. Cell death & disease, 9(10):961, 2018. 9

[62] Congxue Hu, Tengyue Li, Yingqi Xu, Xinxin Zhang, Feng Li, Jing Bai, Jing Chen, Wenqi Jiang, Kaiyue Yang, Qi Ou, et al. Cellmarker 2.0: an updated database of manually curated cell markers in human/mouse and web tools based on scrna-seq data. Nucleic Acids Research, 51 (D1):D870–D876, 2023. 9

[63] Mathias Uhlén, Linn Fagerberg, Björn M Hallström, Cecilia Lindskog, Per Oksvold, Adil Mardinoglu, Åsa Sivertsson, Caroline Kampf, Evelina Sjöstedt, Anna Asplund, et al. Tissue-based map of the human proteome. Science, 347(6220):1260419, 2015. 9

[64] Oana Palasca, Alberto Santos, Christian Stolte, Jan Gorodkin, and Lars Juhl Jensen. Tissues 2.0: an integrative web resource on mammalian tissue expression. Database, 2018:bay003, 2018. 9

[65] Jonathan JH Bray, Rosie Freer, Alex Pitcher, and Rajesh Kharbanda. Family screening for bicuspid aortic valve and aortic dilatation: a meta-analysis. European Heart Journal, page ehad320, 2023. 9

[66] Casey S Greene, Arjun Krishnan, Aaron K Wong, Emanuela Ricciotti, Rene A Zelaya, Daniel S Himmelstein, Ran Zhang, Boris M Hartmann, Elena Zaslavsky, Stuart C Sealfon, et al. Understanding multicellular function and disease with human tissue-specific networks. Nature genetics, 47(6):569–576, 2015. 9

[67] Gregorio Alanis-Lobato, Miguel A Andrade-Navarro, and Martin H Schaefer. Hippie v2. 0: enhancing meaningfulness and reliability of protein–protein interaction networks. Nucleic acids research, page gkw985, 2016. 9

[68] Max Kotlyar, Chiara Pastrello, Nicholas Sheahan, and Igor Jurisica. Integrated interactions database: tissue-specific view of the human and model organism interactomes. Nucleic acids research, 44(D1):D536–D541, 2016. 9

[69] Cui-Xiang Lin, Hong-Dong Li, Chao Deng, Yuanfang Guan, and Jianxin Wang. Tissuenexus: a database of human tissue functional gene networks built with a large compendium of curated rna-seq data. Nucleic acids research, 50(D1):D710–D718, 2022. 9

[70] Junha Cha, Jiwon Yu, Jae-Won Cho, Martin Hemberg, and Insuk Lee. schumannet: a single-cell network analysis platform for the study of cell-type specificity of disease genes. Nucleic acids research, 51(2):e8–e8, 2023.

[71] Daniel V Veres, Dávid M Gyurkó, Benedek Thaler, Kristof Z Szalay, Dávid Fazekas, Tamás Korcsmáros, and Peter Csermely. Comppi: a cellular compartment-specific database for protein– protein interaction network analysis. Nucleic acids research, 43(D1):D485–D493, 2015. 9

[72] Albert-László Barabási and Réka Albert. Emergence of scaling in random networks. science, 286(5439):509–512, 1999. 15

[73] Bryan Perozzi, Vivek Kulkarni, Haochen Chen, and Steven Skiena. Don’t walk, skip! online learning of multi-scale network embeddings. In Proceedings of the 2017 IEEE/ACM International Conference on Advances in Social Networks Analysis and Mining 2017, pages 258–265, 2017. 16

[74] Donna Maglott, Jim Ostell, Kim D Pruitt, and Tatiana Tatusova. Entrez gene: gene-centered information at ncbi. Nucleic acids research, 33(suppl_1):D54–D58, 2005. 18

[75] Chunlei Wu, Adam Mark, and Andrew I Su. Mygene. info: gene annotation query as a service. bioRxiv, page 009332, 2014. 18

[76] David Ochoa, Andrew Hercules, Miguel Carmona, Daniel Suveges, Asier Gonzalez-Uriarte, Cinzia Malangone, Alfredo Miranda, Luca Fumis, Denise Carvalho-Silva, Michaela Spitzer, et al. Open targets platform: supporting systematic drug–target identification and prioritisation. Nucleic acids research, 49(D1):D1302–D1310, 2021. 18

